# Neutralization potential of Covishield vaccinated individuals sera against B.1.617.1

**DOI:** 10.1101/2021.05.12.443645

**Authors:** Pragya D. Yadav, Gajanan N. Sapkal, Priya Abraham, Gururaj Deshpande, Dimpal A Nyayanit, Deepak Y. Patil, Nivedita Gupta, Rima R. Sahay, Anita M. Shete, Sanjay Kumar, Samiran Panda, Balram Bhargava

## Abstract

Covishield comprises the larger proportion in the vaccination program in India. Hence, it is of utmost importance to understand neutralizing capability of vaccine against the B.1.617.1 variant which is considered to responsible for surge of the cases in India. The neutralizing-antibody (NAb) titer against B.1.167.1 and prototype B.1 variant (D614G) was determined of the vaccine sera (4 weeks after second dose) of COVID-19 naïve subjects (n=43) and COVID-19 recovered subjects (n=18). The results demonstrated that sera of COVID-19 recovered subjects (n=18) who received two doses of Covishield have higher NAb response compared to the COVID-19 naive with a significant difference (p<0.0001) in NAb titer against B.1 and B.1.617.1 In-spite of reduction in the neutralizing titer against B.1.617.1 variant; Covishield vaccine-induced antibodies are likely to be protective to limit the severity and mortality of the disease in the vaccinated individuals.

## Text

SARS-CoV-2 variants have emerged and spread to other countries during this pandemic [1]. Many of the studies have reported reduced neutralization capabilities of vaccines viz., mRNA-1273, NVX-CoV2373, BNT162b2, and ChAdOx1-nCoV19 against different variants B.1.1.7, B.1.351 and B.1.1.28 P1 [1-2]. Recently, SARS-CoV-2 variant, B.1.617.1, B.1.617.2 and B.1.617.3 with specific deleterious mutations have been reported from India [3-4], which has been associated with increase in the number of SARS-CoV-2 cases especially in Maharashtra state [3]. This variant has been shown to have higher transmissibility and pathogenicity in hamster model and has raised serious concern pertaining to the national COVID-19 vaccination program in India and other countries [3,5]. Over 177 million doses of vaccine have been administered to Indian citizens with two approved vaccines-Covishield (Astrazeneca-Oxford) and Covaxin™ (BBV152) [6]. Recently, we have demonstrated the neutralizing efficacy of Covaxin™ against B.1.617.1 variant [4,7,8]. Covishield comprises the larger proportion in the vaccination program in India. Hence, it is of utmost importance to understand neutralizing capability of Covishield vaccine against this emerging variant.

We have evaluated neutralizing capability of the Covishield vaccine receipient sera obtained four weeks after the second dose of COVID-19 naïve subjects (n=43) and COVID-19 positive recovered subjects (n=18).

The neutralizing-antibody (NAb) titer against B.1.167.1 and prototype B.1 variant (D614G) was determined for both the categories of sera. Of the sera obtained from COVID-19 naïve subjects, 23.25% samples (n=10) didn’t show NAb titer against both the variants. 27.90% of the samples (n=12) showed NAb titer only with B.1. A total of 21 serum specimens (48.83%) elicited NAb titers against both B.1 and B.1.617.1 variants. The GMT along with standard deviation of Covishield vaccinee sera against B.1 and B.1.617.1 were 42.92±3.8 (95% CI:40.21-128.5; n=43) and 21.92±4.42 (95% CI:24.4-62.64; n=43) respectively.

The results demonstrated that sera of COVID-19 positive recovered subjects (n=18, red color) who received two doses of Covishield have higher antibody response compared to the COVID-19 naive vaccinees (n=43, green color) with a significant difference (p <0.0001) in NAb titer against B.1 and B.1.617.1 variants (Figure 1 A and 1B). An increase in the GMT of the sera of COVID-19 recovered cases with vaccination (29.5 fold) compared to COVID-19 naïve vaccinees (23.5 fold) was observed against and B.1.617.1 respectively. This indicates that COVID-19 recovered cases vaccinated with 2 doses had very high immune response in comparision to COVID-19 naïve vaccinees who received 2 doses.

**Figure 1:**
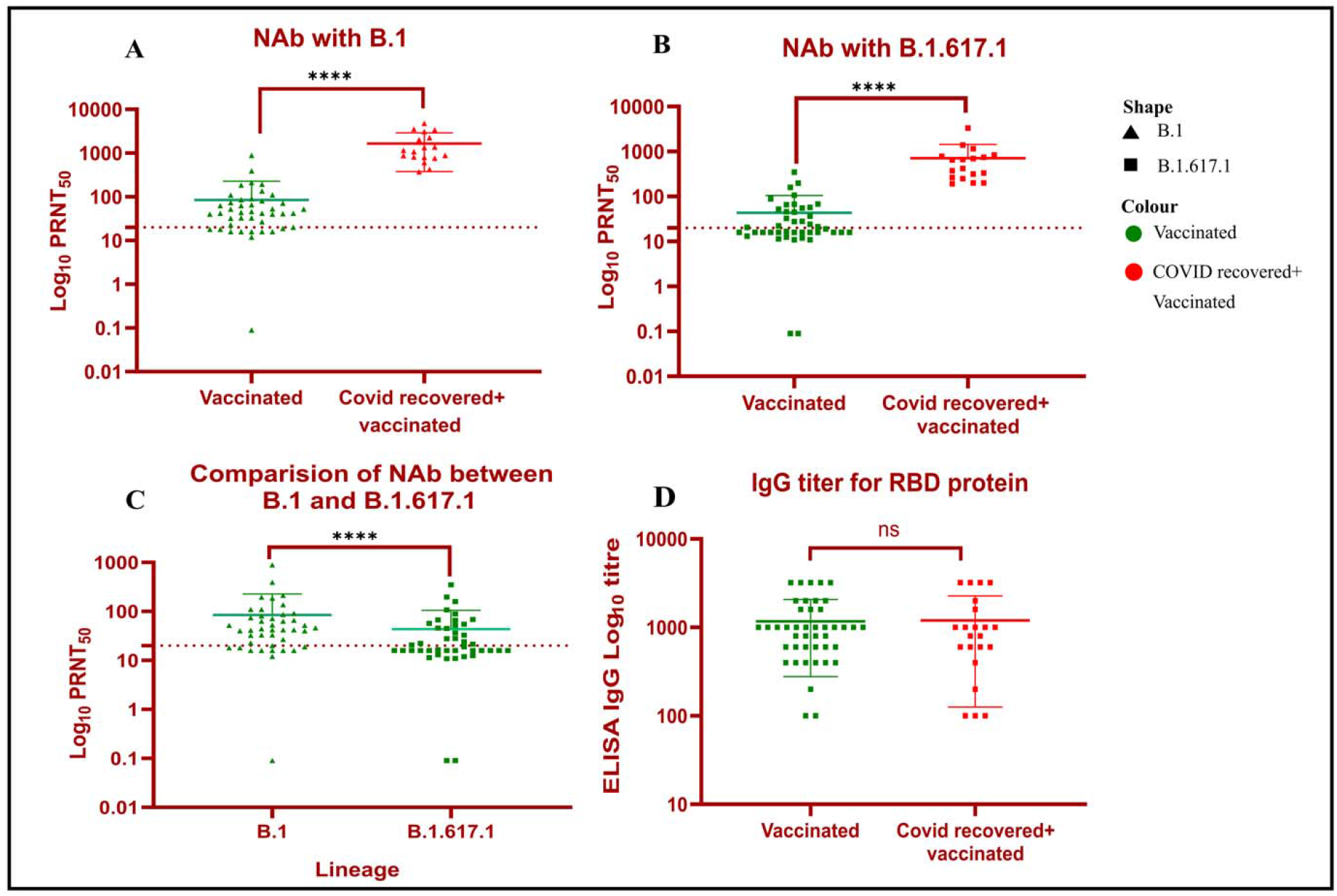
Neutralization of B.1.617.1 variant: Comparison of NAb titer between COVID-19 naïve cases (green, n=43) and COVID-19 recovered cases (red, n=18) administered with 2 doses of vaccine (sera collected 28 days after the second dose) against B.1 **(A)** and B.1.617.1 **(B)**. A two-tailed pair-wise comparison was performed using the Mann-Whitney test and **** represent p-value <0.001 **C)** Scatter plot depicting the neutralization activity of the COVID-19 naïve cases (n=43) vaccinated with two doses of Covishield vaccine (sera collected 28 days after the second dose) against the prototype B.1 (D614G) (green, triangle) and B.1.617.1 (green, square). A neutralization reduction factor of 1.94 was observed between the B.1 (D614G) and B.1.617.1 variant. A two-tailed pair-wise comparison was performed using the Wilcoxon matched-pairs signed-rank test with a p-value of 0.05. **** represent p-value <0.001 and **p value=0.0038,ns= non-significant p-value **(D)** A comparison of IgG antibodytiter against RBD protein between COVID-19 naïve cases (n=43) and COVID-19 recovered cases (n=18) administered with 2 doses of vaccine. A two-tailed pair-wise comparison was performed using the Mann-Whitney test and ns= non-significant p-value.

A pair-wise comparison using Wilcoxon matched-pairs signed-rank test demonstrated a significant two-fold reduction (p-value<0.0001) in the neutralization titer of B.1.617.1 compared to B.1 variant in the COVID-19 naïve vacinees (Figure 1 C). Further, we also determined the IgG titer against S1-RBD and observed a non-significant difference between COVID-19 recovered cases administered with 2 dose of vaccine and COVID-19 naïve vaccinated subjects (Figure 1D).

Inspite of reduction in the neutralizing titer against B.1.617.1 variant; Covishield vaccine-induced antibodies are likely to be protective in limiting the severity of disease and mortality in the vaccinated individuals. Also COVID-19 recovered individuals with immunization can maintain protective antibody titer for longer period.

## Supporting information

Supplementary Information Methodology

## Ethical approval

The study was approved by the Institutional Biosafety Committee and Institutional Human Ethics Committee of ICMR-NIV, Pune, India under the project ‘Propagation of new SARS-CoV-2 variant isolate and characterization in cell culture and animal model’.

## Author Contributions

PDY and PA contributed to study design, data analysis, interpretation and writing and critical review. GNS, GRD, DYP, RRS, AMS, DAN and SK contributed to data collection, interpretation, writing and critical review. NG, SP, and BB contributed to the critical review and finalization of the paper.

## Acknowledgement

Authors gratefully acknowledge the staff of ICMR-NIV, Pune including Dr. Rajlaxmi Jain, Mr. Prasad Sarkale, Mr. Shrikant Baradkar, Ms. AashaSalunkhe and Mr. Chetan Patil for extending excellent technical support.

